# Twig Age 2.0: Adjusting twig age data for differences in palatability among taxa to track ungulate browsing over sites and time

**DOI:** 10.1101/2024.12.06.627192

**Authors:** Donald M. Waller

## Abstract

1. Ungulate browsing threatens forest tree regeneration and diversity across forests in North America and other temperate regions but is inadequately monitored. This reflects both inadequate commitment and the lack of efficient, reliable methods to accurately track these impacts across sites and over time.
2. The twig age method estimates time: how long twigs grow before being browsed, providing a direct measure of herbivory. The original method aged twigs on maple (*Acer*) seedlings, but it works well in other deciduous species. Given ungulate preferences for some species, it is important to adjust twig ages to correct for these.
3. Using multiple datasets from many species and sites across the upper Midwest, I confirm the power and sensitivity of the twig age method for assessing and comparing white-tailed deer impacts over time, among species, and across sites. Twig ages respond differently to seedling heights when deer are present, providing an additional indicator. I extend the original method to rank and adjust for differential browsing among species in two ways: using least square means from statistical models and using ratios to adjust values to an *Acer saccharum* standard. These adjustments match, supporting the use of either to accurately track shifts in browse pressure over time and across sites even when these differ in species composition.
4. The adjusted twig-age method is highly efficient, allowing a sole worker to score 100+ seedlings/saplings within an hour to accurately gauge real-time ungulate impacts at a site. The reliability, power, and versatility of adjusted twig age data set a standard for assessing browse pressure and provide an efficient vehicle to expand programs to monitor deer impacts across landscapes.

---

> “A common error is to try to appraise [deer numbers] by census, rather than by browse conditions. The public can dispute endlessly about censuses, but it cannot dispute dead browse plants.” Aldo Leopold, 1947

Leopold recognized in the 1930s and 40s that high ungulate densities could impair plant cover, diversity, soil stability, and other habitat values. His prescient efforts to alert and inform others about these concerns, however, were largely ignored even as deer populations and impacts increased further through the latter 20^th^ century (Alverson et al., 1988; F. L. Russell et al., 2001; Augustine & DeCalesta, 2003; Côté et al., 2004). Deer now act as a keystone herbivore, greatly altering species composition, abundance, and ecosystem processes in many temperate forests (McShea & Rappole, 1992; Waller & Alverson, 1997). Most immediately, abundant ungulates (henceforth deer) feed heavily on favored species, including many wildflowers (e.g., Augustine & Frelich, 1998), palatable woody shrubs, and juvenile trees. This browsing has substantially reduced the density and diversity of many plants in forest understories while enhancing populations of grasses, sedges, and invasive species (Rooney et al., 2004; Gorchov et al., 2021). Where deer densities remain well above historical levels, ‘recalcitrant understories’ of ferns and tough shrubs have replaced diverse wildflower communities, limiting tree regeneration (Royo & Carson, 2006). Browsing on oaks (*Quercus*) and slow-growing, shade-tolerant conifers like eastern and western hemlock (*Tsuga*), norther white and western red cedar (*Thuja*), and Canada yew (*Taxus*) in North America has made these scarce as well, creating shifts in forest composition that persist for decades (Allison, 1990; Stange & Shea, 1998; Alverson et al., 1999; Rossell et al., 2005; Sabo et al., 2018; Patton et al., 2021; Webster & VanderMolen, 2023). Parts of the Allegheny plateau of Pennsylvania are now overbrowsed “fern parks” that may convert to savanna (Wigg et al., 2011). Deer browsing has thus created a substantial “regeneration debt” across much of northeastern North America (Bradshaw & Waller, 2016; Miller & McGill, 2019).

To protect sensitive tree seedlings from deer, foresters now commonly fence or place tubes over individual seedlings, greatly increasing costs (Beguin et al., 2016). High deer densities also appear related to the spread of invasive earthworms (Dobson & Blossey, 2015), declines in songbird species (Chollet & Martin, 2013; Tymkiw et al. 2013), vehicular accidents (Raynor et al., 2021), and increases in wildlife and human diseases including Chronic Wasting Disease in ungulates and tick-borne human zoonoses including Lyme disease, Ehrlichiosis, tick fever, etc. (Wiznia et al., 2013; Kilpatrick et al., 2014; Larson et al., 2022).

Given all these impacts, it behooves forest and wildlife managers to combine forces to monitor deer browse impacts closely and continuously over landscapes they oversee. To date, wildlife managers have mostly focused on estimating deer densities and carrying capacities (K) with the goal of sustaining deer populations near their estimated maximum sustained yield. This has hardly limited deer impacts, particularly as hunting continues to lose effectiveness in many areas. This has accentuated controversies surrounding deer management (Martin et al., 2020; Blossey et al., 2024). In addition, estimating deer densities is expensive, time consuming, and controversial and generates numbers that often fail to predict deer impacts because they are too broad in scale and because plant populations often express delayed and sustained responses to herbivory (Augustine et al., 1998; Webster & VanderMolen, 2023). It may often be simpler, cheaper, and more relevant to measure deer impacts directly (Sabatini et al., 2023).

Methods to estimate deer impacts include fenced exclosures (Allison, 1990; Kain et al., 2011; Abrams & Johnson, 2012; Flagel et al., 2016); natural refugia like islands, boulders, or cliffs (Rooney & Dress, 1997; Mudrak et al., 2009); differential shifts in understory composition (Rooney et al., 2004; Begley-Miller et al., 2014); the abundance and flowering condition of indicator taxa (Balgooyen & Waller, 1995); rates of browsing on extant stems (Morellet et al., 2007); rates and ratios of browsing on particular species (Frerker et al. 2013); and the density and/or height of stump sprouts (Royo et al. 2016), naturally occurring woody seedlings (Hett & Loucks, 1976; McWilliams et al., 2012; McWilliams et al., 2015; Blossey et al. 2017; Quirion & Blossey, 2023) or planted “sentinel” seedlings (Blossey et al., 2017). Each of these methods has advantages and disadvantages, but note that seedling counts and heights are often affected by local conditions, reducing their reliability, while transplanting ‘sentinel’ seedlings demands much time and effort.. Reassuringly, a 25-year study in the Italian alps found browsing ratios on conifers reliably tracked changes in ungulate density (Donini et al. 2024). Less reassuringly, Patton et al. (2018) found data on deer-vehicle collisions and Lyme disease incidence predict deer browse impacts more reliably than forest regeneration metrics based on seedling numbers (an extension to the USFS FIA forest inventory program). Reviewing methods exceeds the scope of this paper, but continuing uncertainty about which methods and data to use strongly suggests a need for additional tests and comparisons before we can determine which indicators are most accurate, reliable, and cost-effective, i.e., a solid foundation for efficient, standardized monitoring programs.

Here, I assemble twig age data from the upper Midwest, USA, to assess how efficient, versatile, and reliable the twig age method is and how it can be adjusted to generate more accurate and reliable comparative data. Rather than relying on the seedling density, height, size, or the fraction of stems browsed, the twig age method was designed to measure the frequency at which deciduous available twigs are browsed (Waller et al., 2017). Specifically, it seeks to estimate how long a twig can grow unmolested before being bitten off by visually aging twigs using terminal bud scale scars. Once learned, the method is quick to apply, allowing many measurements in short period of time with minimal equipment (Waller, 2018). Goals here are to:

1. confirm that twig ages respond sensitively to differences in deer browse intensity;
2. compare how twig ages compare to seedling heights for indicating browsing;
3. assess how well twig ages track changes in browsing over time at one site;
4. examine changes in twig ages with seedling height among species to determine whether these patterns themselves reflect browse impacts;
5. assess how well twig ages serve to compare browsing intensities over species and sites;
6. examine how independent (additive) species and site effects are to confirm that these differences are consistent (transitive);
7. if deer impacts are transitive, explore how to correct for different species’ sensitivity to browsing to obtain a standardized metric to reliably compare browsing impacts among sites.

## Methods

### The twig Age Method

The twig age method relies on closely inspecting twigs in the ‘molar zone’ of ungulates to estimate their age by counting terminal bud scale scars (Waller et al. 2017). It thus requires knowledge of woody species and some training or experience to identifying these features. Tools, however, are simple: a measuring tape (to measure heights), a map or GPS device (to locate sites), and a method to record data (e.g., audio recorder or clipboard/notebook). At each study site, the observer decides on a set of woody deciduous species to sample and then moves through the site to locate seedlings and saplings (henceforth “seedlings”) of those species. Each plant is then sampled by recording its height and aging two live twigs located between 20-180cm above the ground (on different branches). Aging twigs involves working in from the terminal part with this year’s leaves to the “parent” branch that generated that twig, counting terminal bud scale scars to age the daughter twig. First-year twigs with leaves attached directly to their parent branch are one year old. Twigs with one terminal bud scale scar before the parent branch are scored as 2-year old twigs, etc. Counts are halted at 5 years as older twig are rare and hard to age accurately. Parent branch ages are ignored. The method may also record the presence of fresh browsing on this year’s twigs. Data for each seedling thus consist of date, species sampled, height, two twig ages, and whether twigs show evidence of recent browsing. Site data include location, observations on canopy composition or forest type, and start and stop times.

### Study Sites

I report results from three regions sampled in different years. The twig age method was originally based on 2015-1027 results for three maples (*Acer rubrum, saccharum,* and *pensylvanicum*) under an old-growth canopy at the Huron Mountain Club (HMC, Fig. 1, SI Table 1). Revisits to this site in 2019 and 2022 generated more data for these maples plus data for *Betula alleghaniensis*, *Ostrya virginiana*, and *Quercus rubra* in 2022. This site includes a 2 ha fenced deer exclosure, meaning samples from in- and outside the fence allow us to rigorously assess effects of deer browsing. This remote site had limited access, meaning breaches of the fence from tree and limb falls were only repaired during the growing season. Deer entered the exclosure in some winters, meaning estimates of deer impacts based on twig ages differences across the fence underestimate actual deer impacts.

**Figure 1.**
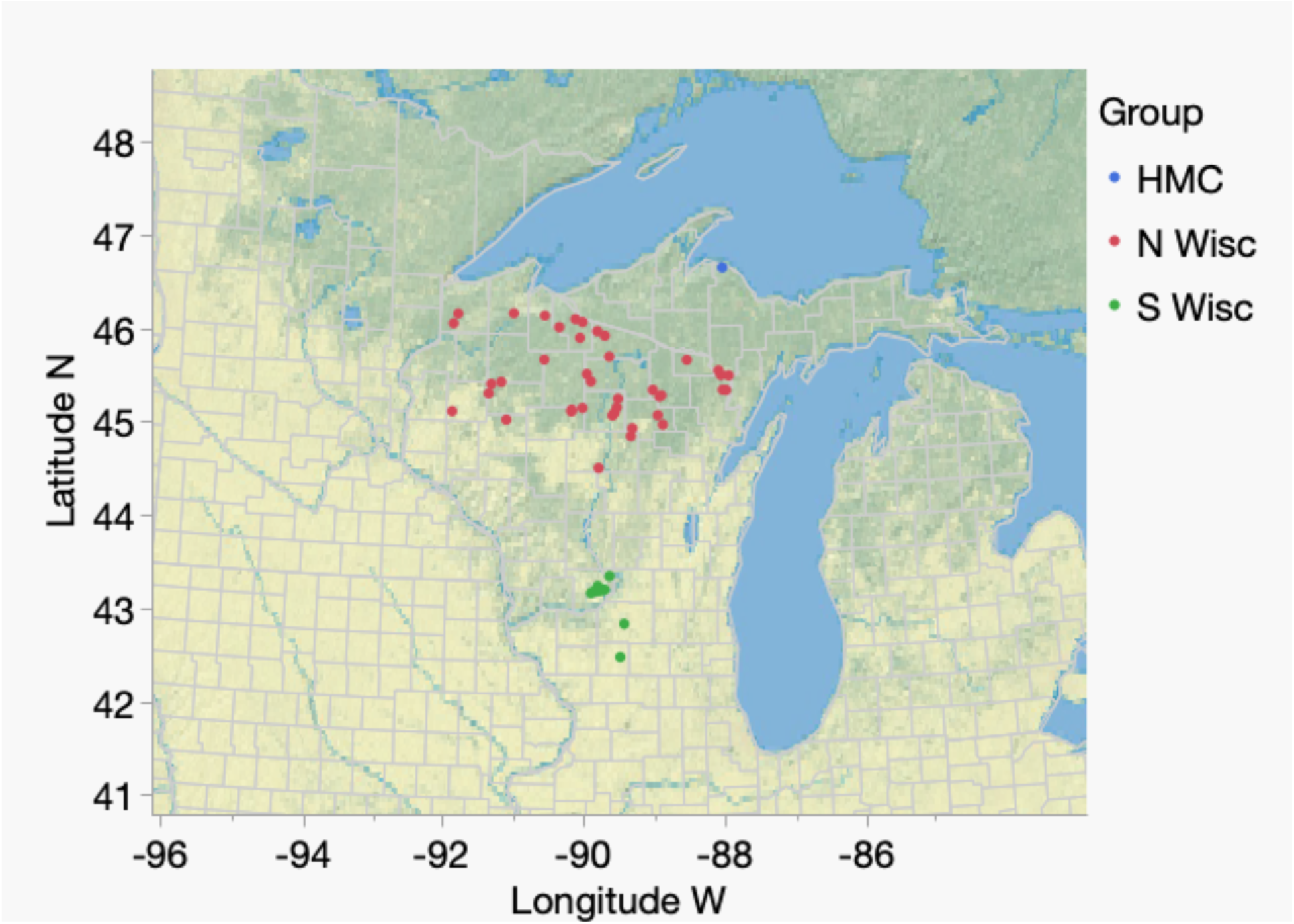
Map of the upper Midwest showing locations of the 50 study sites. These include the Huron Mountain Club (HMC) in the Upper Peninsula of Michigan, nine sites in scattered mesic forests in southern Wisconsin, and 40 upland sites in the more continuously distributed forests of northern Wisconsin.

In May-July, 2019, I aged twigs of up to eight taxa (*Acer rubrum, Acer saccharum, Carya cordiformis, Carya ovata, Fraxinus* spp, *Quercus alba*, *Quercus bicolor*, and *Quercus rubra*) at nine sites in southern Wisconsin (Fig. 1, SI Table 1). These occur within medium to large forest fragments located within mostly agricultural landscapes. I strove to sample seedlings in rough proportion to their abundance.

In summer, 2023, I measured twig ages at 40 sites across northern Wisconsin (Fig. 1, SI Table 1), again sampling species as they were encountered. Sites occurred mostly on County Forest lands with young to medium-aged canopies chosen for another study (Brice et al., in prep). I sampled common shrubs at a few sites where tree seedlings were scarce. I omit analyses of rarer taxa (<56 seedlings) that occurred at only a few sites (*Alnus, Amelanchier, Carpinus, Carya, Rhus*, and *Ulmus*), focusing instead on *Betula, Acer, Cornus, Fraxinus, Ostrya, Populus, Prunus, Quercus*, *Tilia,* and *Viburnum*. Species were merged to genus except in *Acer* and *Quercus*.

### Analyses

These simple data permit multiple approaches and alternative indicators. Individual woody seedlings are the sample unit, so I first averaged the twig ages within seedlings to obtain reliable results. Distributions of these mean twig ages confirmed that values vary over species and sites. The twig-age metric is statistically well-behaved, meeting assumptions of general linear models (e.g., few/no outliers; linear relationships; Normally distributed residuals, etc.). In applying GLM’s, I checked Goodness of Fit statistics and interactions before settling on final, usually simpler, models. Taxa, exclosure status, and site were analyzed as fixed effects as these variables are all of prime interest. For clarity and simplicity, I present illustrative rather than exhaustive analyses, often based on subsets of the data.

The fenced exclosure at HMC allowed explicitly tests of deer effects on seedling heights and twig ages in maple (*Acer*) and other species. I first analyzed within year variation to assess effects of species, seedling height, and exclosure (deer), plus their 2-way interactions. I then pooled the data to analyze effects of year in combination with other predictors to assess shifts in deer effects over time. Finally, I used data on three more species in 2022 to evaluate how twig ages differ among species.

Data for multiple taxa across many sites in southern and northern Wisconsin allowed me to explore their differential sensitivity to deer impacts, how consistent these differences are across sites, and how twig ages and inferred deer impacts can be compared among sites. These data represent “natural” factorial experiments, allowing me to use two-way ANOVA’s to assess separate and combined effects of taxa sampled and site on twig age variation. By adjusting site for species effects and vice versa, these analyses provide more precise and less biased estimates of effects. The interaction between these factors tests whether species and sites vary consistently in twig age and inferred deer impacts. This independence was supported in the S Wisconsin results (site x species interaction F=1.89, p=0.08, N=698) but not in N Wisconsin (interaction F=2.07, p<0.001, N=4110). Still, even this significant interaction is small relative to the main effects of site (F=20.8) and taxon (F=24.6), suggesting these can predict twig age effects with good power (r^2^=24.1%). The consistent ways in which twig ages responded to species and site effects (see Results) reinforced the idea that this metric provides a reliable method to rank and compare species and sites.

Sample sizes ranged from 32 to 303 (mean: 120.8, SI Table 1) seedlings per site. All data were analyzed using JMP Pro Vers. 17 or 18 (SAS Institute, Cary, NC).

### Adjusting twig ages to compare sites

The 2-way ANOVAs generate adjusted least-square (LS) means for main effects, adjusting both species and site effects for the other variable, reducing bias. These LS twig age means (Adjusted Twig Age 1, or ATA1) provide a way to estimate site effects while controlling for differences among sites in how species composition affects twig age (i.e., how palatability differs among species). A second approach to adjust for differences in palatability among species is to recompute twig ages within species using a universal standard, e.g., sugar maple (*Acer saccharum*). Sugar maple (SM) is the most abundant and widespread species at sites across both S and N Wisconsin. Sugar maple can also resprout after browsing and has lower palatability than many other species, allowing it to persist at most sites even under moderately strong browsing. Because twig ages vary consistently between species, I adjusted each species’ twig age to the SM standard by calculating relative palatability: the ratio in mean twig age at each site, ***j,*** between SM and all other taxa, ***i,*** present (using only sites with at least 4 SM seedlings):

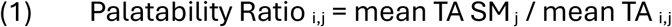

This estimates the palatability of each species relative to SM. A weighted average of these palatability ratios across sites, weighted by local sample sizes for that species (***N i,j***) generated a mean palatability ratio for each species:

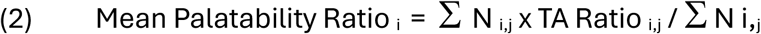

A 2-way ANOVA showed linear, additive effects on this ratio (i.e., no significant species x site effect, F=0.84, p=0.58, N=171). This consistency means SM adjusted palatability ratios can be combined across sites to compute standardized twig age values, using all data available to enhance sample sizes and more precisely compare sites that share few species. I used these palatability ratios to adjust twig ages to the common SM standard by multiplying each species’ TA at a site by its mean palatability ratio. Averaging these SM adjusted values over all species present (weighting by abundance) generates a second adjusted estimator (ATA2).

Finally, I calculated two other summary variables: first, a site’s Sugar Maple Browse Index, defined as the converse of ATA2:

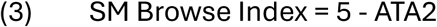

This browse index increases as browse pressure increases, providing intuitive values for forest and wildlife managers. Second, a site’s Community Palatability score is its mean palatability ratio (Eqn. 2 above), weighted by species’ abundances.

To assess consistency, I compared the four twig age metrics (simple mean twig age, local SM TA, ATA1, and ATA2) using linear regressions. I next regressed site Community Palatability scores against the four twig age estimators to test whether communities showing more current browse (higher twig ages) also display lower palatability. Such a result would suggest that more heavily browsed site are shifting in species composition toward less palatable species.

Finally, I mapped browse intensity across the 40 sites in N Wisconsin and tested for latitudinal and longitudinal trends. Such maps further allow us to assess spatial patterns in deer browse and generate hypotheses regarding their causes.

## Results

### Efficiency

In 2023, I used a cell phone tucked into a shirt pocket to continuously record spoken data, leaving both hands free to measure heights and inspect and age twigs. This greatly accelerated data collection relative to writing data. I obtained good samples at most sites within an hour (mean: 44 min., mean rate: 2.4 seedlings/min). Additional time was then needed to enter the data into spreadsheets, but similar times are needed to enter written data.

### Height vs. twig age indicators

The exclosure at HMC allowed us to assess direct effects of deer on both seedling height and twig growth among *Acer* species in 2015, 2016, and 2017 (cf. Waller et al. 2017). Later 2019 results allow further comparisons (Table 1, Fig. S1). Both indicators show significant effects of deer, *Acer* species, and their interaction, but deer affect twig ages far more than height (R^2^ = 0.73 vs. 0.41 and F = 612 vs. 6.2, respectively). *Acer pensylvanica* showed a particularly striking response with seedlings growing slightly taller, on average, inside the fence (117 vs. 98.4cm) but with much higher twig ages (2.32 vs. 1.14 yrs). Twig ages are also less sensitive to the species being measured with a much smaller deer x species interaction effect (F=3.9 vs. 64.3, Table 2, Fig. S1). Twig ages also responded differently to deer, showing declines with seedling height outside the fence but not inside (see also *Height effects* below). This suggests deer browsed more on taller seedlings at HMC. Independent data from 2022 at HMC show that deer reduced mean twig age by 0.783 years and account for 21.3% of the total variance in twig age vs. 2.0% for height (Fig. 2, SI Table 2). This effect was consistent across species (species x exclosure F=2.06, p=0.07). The equivalent analysis for seedling height shows that deer reduced height by 5.7cm with a model R^2^=0.19 (vs. 0.52) and an Exclosure F=6.38 (vs. 183.3). I also found no correlation between mean seedling heights and mean twig ages among the 40 N Wisconsin sites (r^2^=0.0007, p=0.87). Thus, height is not measuring what twig age is measuring. In sum, twig age has high power to detect and track deer impacts whereas seedling heights respond far less to deer and far more to other factors.

**Figure 2.**
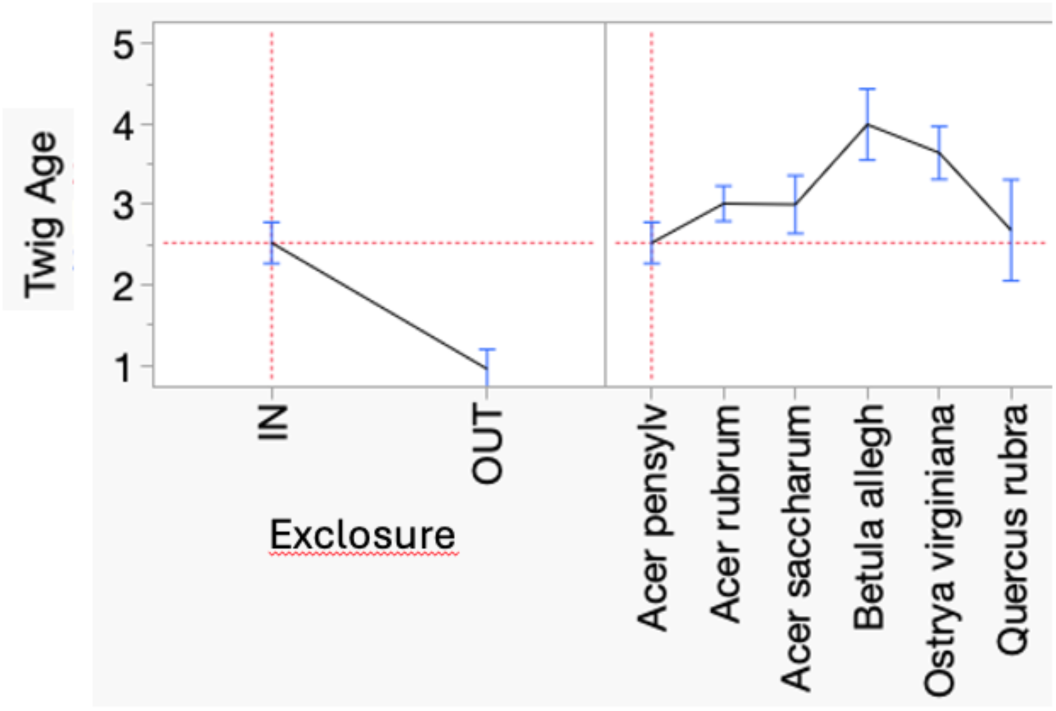
Variation in twig age measured over six species at HMC in 2022. Least square means and standard errors are plotted, adjusted for other effects. The exclosure provides a lower bound estimate for deer effects. The species x exclosure interaction was not significant (see GLM analysis, SI Table 2). Overall r2=0.52, N=247.

**Table 1.** General linear models analyzing variation in a) seedling height and b) twig age among 288 seedlings of three maple species sampled at HMC in 2019. The twig age metric is more sensitive to deer, less sensitive to species, and has higher explanatory power (R^2^ = 0.73 vs. 0.41).

| Source | Nparm | DF | Sum of Squares | F Ratio | Prob > F |
| --- | --- | --- | --- | --- | --- |
| Deer Exclosure | 1 | 1 | 6565.13 | 6.2236 | 0.0132* |
| Species | 2 | 2 | 135560.03 | 64.2545 | <.0001* |
| Species x Deer | 2 | 2 | 42968.35 | 20.3667 | <.0001* |

b) Twig age
| Source | Nparm | DF | Sum of Squares | F Ratio | Prob > F |
| --- | --- | --- | --- | --- | --- |
| Deer Exclosure | 1 | 1 | 341.47 | 611.9307 | <.0001* |
| Species | 2 | 2 | 4.33 | 3.8797 | 0.0218* |
| Species x Deer | 2 | 2 | 26.65 | 23.8748 | <.0001* |

**Table 2.** HMC. General linear model of variation in twig age from 2015 to 2019 at HMC as a function of four predictor variables. Only significant interactive effects are shown. Overall r^2^=0.56, N=880.

| Source | DF | SS | F Ratio | Prob > F |
| --- | --- | --- | --- | --- |
| Exclosure | 1 | 624.80 | 796.46 | <.0001* |
| Species | 2 | 68.99 | 43.97 | <.0001* |
| Year | 1 | 39.01 | 49.73 | <.0001* |
| Exclosure*Year | 1 | 41.15 | 52.46 | <.0001* |
| Species*Year | 2 | 14.90 | 9.50 | <.0001* |
| Species*Exclosure | 2 | 4.88 | 3.11 | 0.0451* |
| Height | 1 | 6.15 | 7.83 | 0.0052* |

### Tracking browse impacts over time

Using HMC maple twig age data from 2015 to 2019, a full GLM shows that twig ages respond sensitively to changes in browsing intensity over time (Table 2, SI Fig. S1). In fact, Year has the second largest effect after Exclosure generating responses as both as a main (additive) effect and via its interaction with deer. Thus, deer impacts changed at HMC by increasing recentky. Mean twig ages also varied among species with these effects depending on year somewhat (Table 2). Thus, twig ages respond sensitively to changes in deer browsing over time (Year effect and Year*Exclosure interaction) and by shifting how species respond (Species*Year interaction).

### Browse impacts vary over species and sites

Because twig ages respond strongly to deer (exclosure) effects and how these change over time, we can conclude that twig ages generate a useful metric for tracking “real-time” changes in browse pressure. Although twig ages responded sensitively to taxon at HMC (Fig. 2, SI Table 2), they respond similarly to deer browsing across species (i.e., the species x exclosure interaction lacked significance). The far more extensive S and N Wisconsin surveys confirm that twig ages vary greatly over both species and sites (Fig. 3, Table SI 3). Differences among taxa spanned a 1.6-fold range of twig ages in S Wisconsin in 2019 (Fig. 3a) and 2.3-fold range in N Wisconsin in 2023 (Fig. 3c). Differences among sites were just as pronounced covering 2.3-fold (Fig. 3b) and 2.4-fold ranges (Fig. 3d). These differences among species and sites spanned 1-2 yrs in mean twig age relative to standard errors of 0.2 yrs or less. Twig ages thus emerge as a sensitive tool for comparing and ranking deer impacts among species and sites.

**Figure 3.**
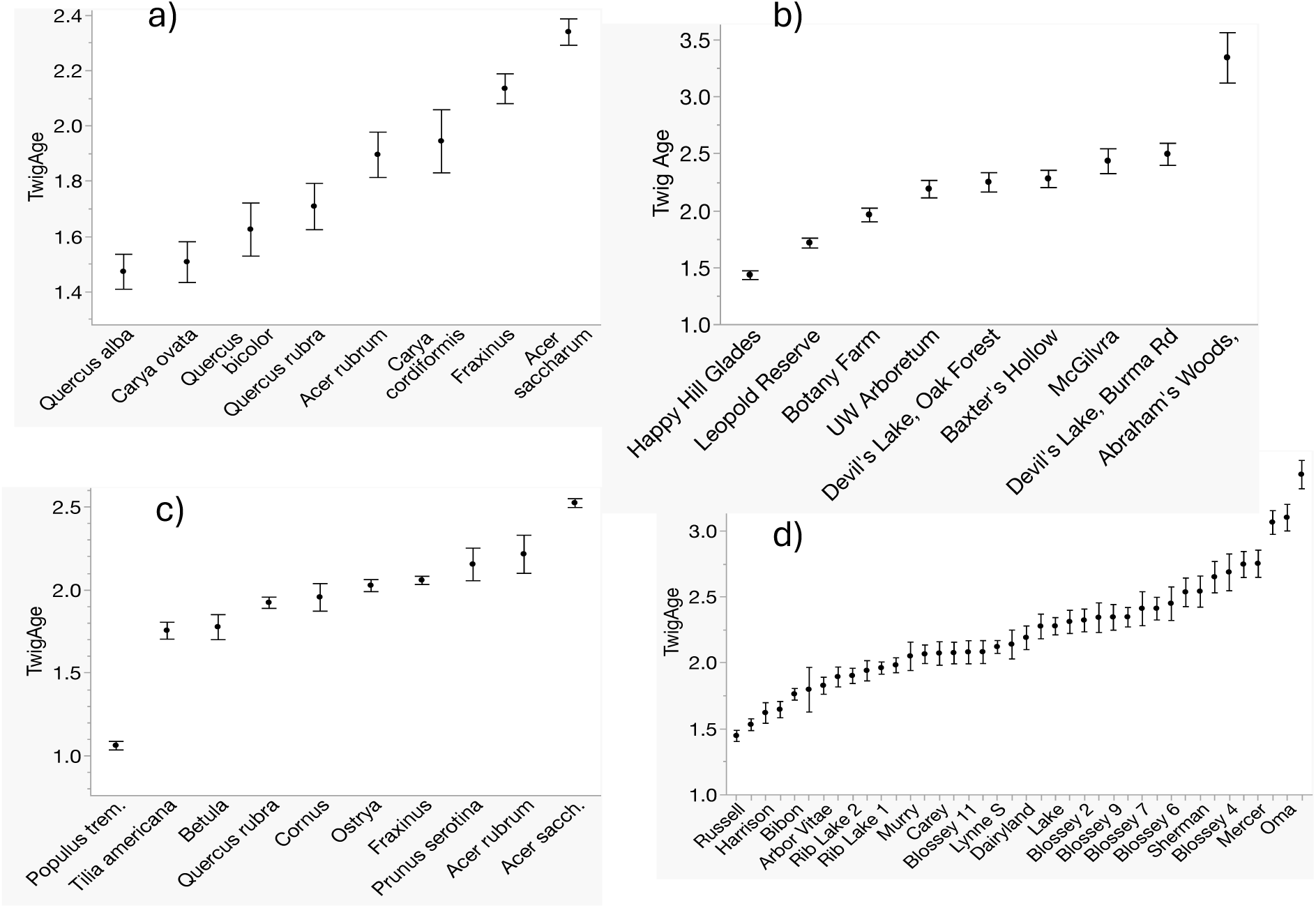
Variation in mean twig ages (unadjusted) among taxa (a,c) and sites (b,d) across the nine S Wisconsin (a,b) and 40 N Wisconsin sites (c,d). Error bars represent 1 SE. GLM analyses of these data appear in SI Table 3.

### Are twig age estimates of browse impacts consistent?

Species display parallel patterns in how their mean twig ages differ, both among sites within a region and between S and N Wisconsin. Aspen (*Populus tremuloides*), birch (*Betula*), hickory (*Carya*), and oak (*Quercus*) generally show high sensitivity to browsing with younger twig ages while the generally older twig ages in sugar maple (*Acer saccharum*) suggest it is less palatable (Fig. 3a, c). This consistency across species suggests that browsing preferences remain consistent enough to rank species in a transitive order. This implies that differences in mean twig age among sites are consistent enough to rank and compare relative browse intensity (Fig. 3b, d). Because these plotted mean site values have not been adjusted for the effects of species, however, some of the apparent differences among sites could reflect differences in species composition. Such artefacts could bias comparisons of browsing pressure among sites particularly if browsing tends to eliminate browse-sensitive species from heavily browsed sites, skewing mean twig ages higher. This appears to be the case as adjusted least square mean twig ages (ATA1) are lower than raw values (SI Fig. S3a). This effect suggests that we can adjust for potential bias by using least square twig age means from 2-way ANOVAs adjusted for species effects (ATA1). Reassuringly, correcting for species effects in this way hardly affects the ranking of sites for browse impacts relative to raw values (rank correlation: ρ=0.91, SI Fig. S3b).

### Height affects twig age

Twig ages in all three regions varied among seedlings of different height, often in a species-specific way. Protected from deer, twig ages tended to increase with height in *Acer pennsylvanicus*, *Ostrya virgininiana*, and *Betula allegheniensis* at HMC but decreased with height in *Acer rubrum*, *A. saccharum*, and *Quercus rubra* (SI Fig. S4). These trends presumably reflect inherent propensities for twigs to live longer or shorter lives as seedlings grew taller. Outside the fence, however, twig ages either remained low or declined on taller seedlings. Across many sites in S and N Wisconsin, it was unprotected seedlings of intermediate height that showed reduced twig age (Fig. 4). This “bite out of the middle” pattern likely reflects how deer prefer to browse seedlings of intermediate height (i.e., an optimal foraging reponse). Twigs on very short or fast-growing tall seedlings thus appear to survive longer. Over all species, these quadratic effects were highly significant in both S and N Wisconsin (SI Table S4). They were also individually significant within *Acer* (F=4.0), *Carya* (F=3.6), and Quercus (F=5.0) in S Wisconsin (all p<0.001).

**Figure 4.**
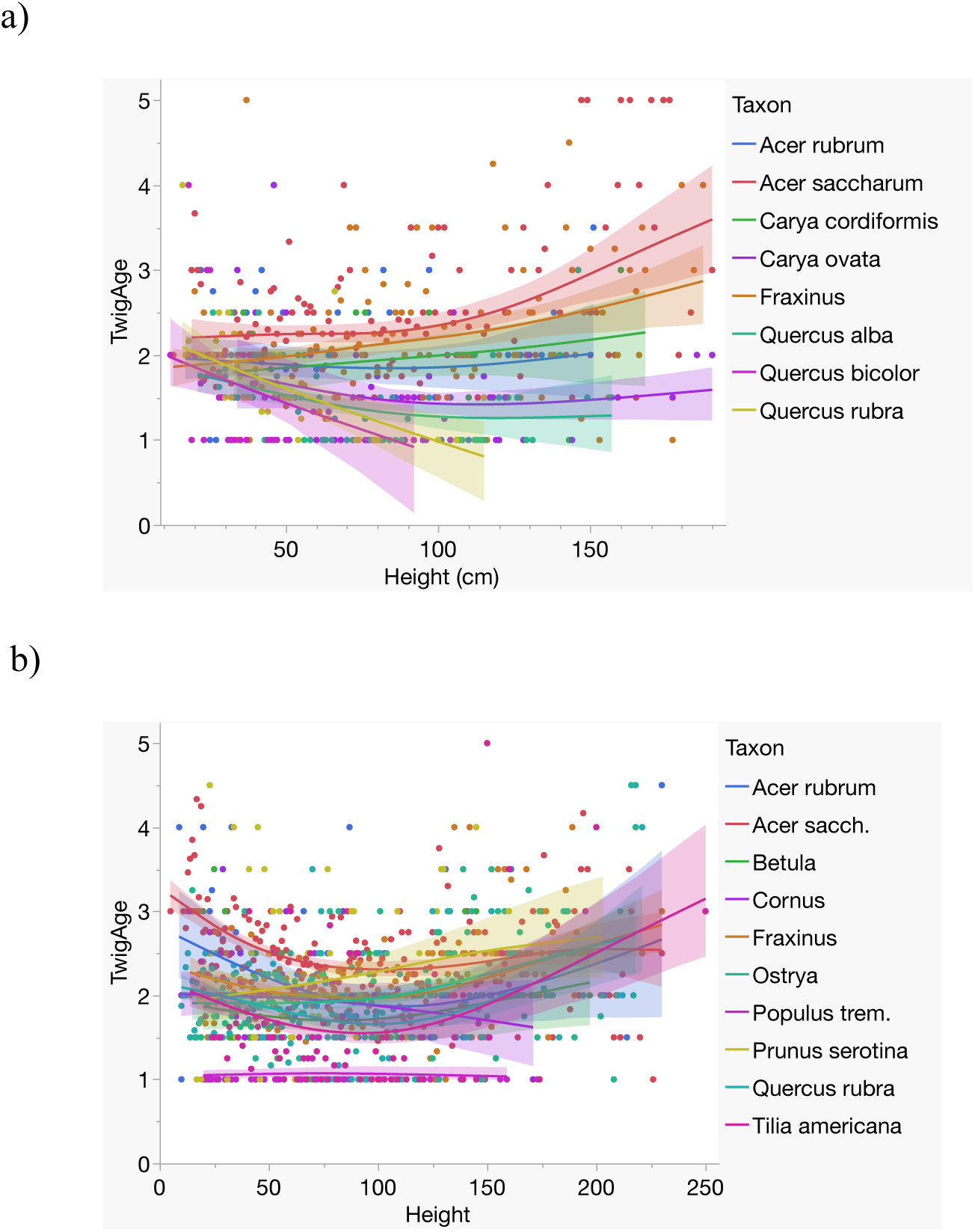
Twig age plotted against seedling height in woody taxa at 9 S Wisconsin sites (a) and 40 N Wisconsin sites (b). In a), the taxa are from the top: Acer saccharum, Fraxinus, Carya cordiformis, Acer rubrum, Carya ovata, Quercus alba, Q. rubra, and Q. bicolor. Both S and N Wisconsin results show strong quadratic effects (SI Table 4).

### Standardizing twig age data

Twig ages of other taxa covary positively with mean sugar maple twig age (SM TA) at the same site (r=0.24, p=0.01, N=176, N Wisconsin), suggesting common responses to deer. In addition, correlations in *Fraxinus*, *Cornus*, *Ostrya*, and *Acer rubrum* were all individually significant. A GLM predicting twig age across the 40 sites reveals that SM TA strongly affects twig ages in other species (F=20.1, p<0.0001, r^2^=0.32, N=171) with linear, additive effects (no interaction, SI Table S5). Given this consistency in how species respond to deer over sites, one can adjust species’ twig age values to a common standard using ratios. I used ratios of mean sugar maple twig age to mean twig ages in other taxa to compute adjustment factors, weighting by taxon abundance (see *Methods*). Local ratios show variation, as expected (SI Fig. S4), but taxon accounts for more than half the variation in these weighted mean ratios (r^2^=0.52, F=19.53, p<0.0001) generating small standard errors (2.4-12.3%, Table 3). These ratios range from 1 (SM) to 2.61 (aspen), revealing deer preferences mirroring those in Fig. 3c. Using these ratios, I adjusted mean twig ages for each species at each site to SM-equivalent values, again weighting by abundance, to generate a standardized SM-equivalent twig age (ATA2) for each site. These values covary with other twig age measures: mean SM TA (0.77), raw mean twig age (all species, r=0.87), and especially the least-square means (ATA2, r=0.97, SI Figure S6). Thus, all measures quantify deer impacts and the last two are interchangeable. This equivalence reflects how the ATA1 and ATA2 indicators both use twig age data for all species while correcting (in different ways) for differential deer impacts on different species.

**Table 3.** Sugar Maple adjustment ratios computed as an average of the ratio of SM to other taxa twig age across sites where the species co-occurred, weighted by taxon abundance at those sites. Ratios represent multipliers to convert twig ages from other taxa to the SM standard. On average, *Populus tremuloides* was most sensitive to deer browse and sugar maple the least. Std Error values use a pooled estimate of error variance.

| **Taxon** | **N** | **Sugar maple**<br>**adjustment ratio** | **S.E. adjustment**<br>**ratio** |
| --- | --- | --- | --- |
| *Populus tremuloides* | 56 | 2.61 | 0.1264 |
| *Tilia americana* | 250 | 1.48 | 0.0598 |
| *Betula* | 66 | 1.44 | 0.1164 |
| *Viburnum* | 38 | 1.34 | 0.1535 |
| *Quercus rubra* | 549 | 1.32 | 0.0404 |
| *Ostrya* | 396 | 1.28 | 0.0475 |
| *Acer rubrum* | 90 | 1.24 | 0.0997 |
| *Fraxinus* | 959 | 1.16 | 0.0306 |
| *Cornus* | 44 | 1.13 | 0.1426 |
| *Prunus serotina* | 56 | 1.06 | 0.1264 |
| *Acer saccharum* | 1500 | 1.00 | 0.0244 |

### Statistical precision

How many seedlings suffice to estimate twig ages reliably? Larger sample sizes increase how precisely we can estimate means as evident in the equation: SE=S.D./√N. This effect was conspicuous for sugar maple estimates in N Wisconsin (Fig. 5a) as well as mean TA and ATA1 (Fig. 5b). Samples of 30-50 seedlings suffice to obtain fairly accurate estimates of sugar maple twig age (SE<0.1). Samples of 100-120 reduce the standard error of overall twig age (raw or adjusted for species effects) to 0.05-0.10 years, small enough to sensitively test among-site differences (Fig. 3b, d). Such sample sizes thus reflect a convenient trade-off between effort and precision for detecting differences in browsing impacts among sites.

**Figure 5.**
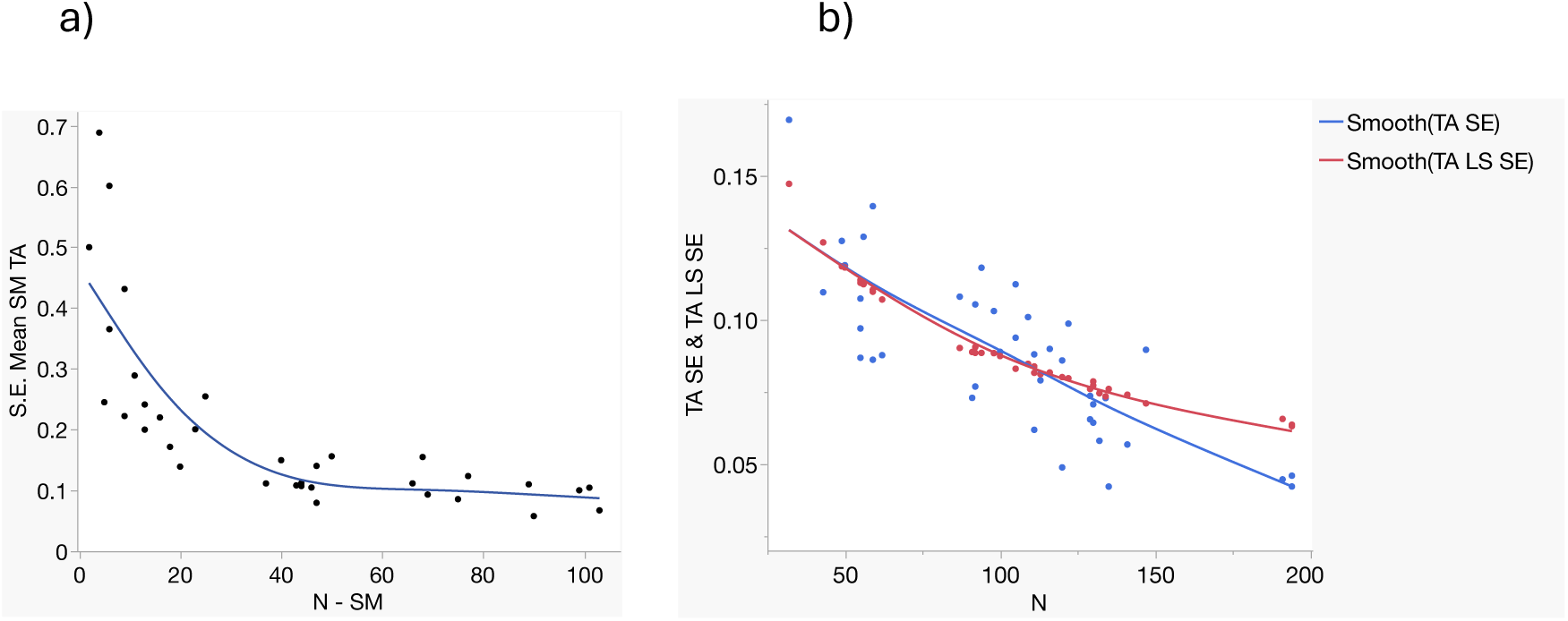
Standard errors (SEs) of estimates for twig age decline as sample sizes (N) increase for data from N Wisconsin. a) Errors for sugar maple twig age (SM TA) estimates decline rapidly until sample sizes exceed 50 seedlings. B) Errors for raw mean twig age (blue, all species) and least square mean twig age (ATA2, in red, adjusted for species effects) have similar declines until samples exceed 130 seedlings.

### Has browsing changed community composition?

Regressing overall site palatability against four estimates of twig age revealed only one significant decline for raw mean twig age (r=-0.47, p=0.0025, SI Figure S7). This is puzzling if all estimate the same thing, but recall that our rationale for adjusting raw twig age for species effects (ATA1 and ATA2) was to obtain more accurate and reliable indicators. The fact that these improved indicators show no relation to palatability suggests that the decline with raw twig age simply reflects how twig ages increase in less palatable species. Wherever these are common, site palatability necessarily falls. This result reinforces the greater reliability of using ATA1 and ATA2. Sherman Corners had the lowest palatability (0.9) where field notes confirmed low cover, few seedlings, and heavy fresh browsing.

### Geographic variation

Neither latitude nor longitude affected mean twig age. However, mapping browse data from the 40 N Wisconsin sites reveals how deer browse intensity varies across these landscapes (Fig. 6). Such maps have value for graphically illustrating the extent and severity of browse threats to regenerating trees and stimulating ideas on what factors might be responsible. Mapping changes in browse intensity over time might also reveal how browse impacts respond to changes in weather and deer and forest management. Causes for patterns in Fig. 6 deserve further attention but exceed the scope of this paper.

**Figure 6.**
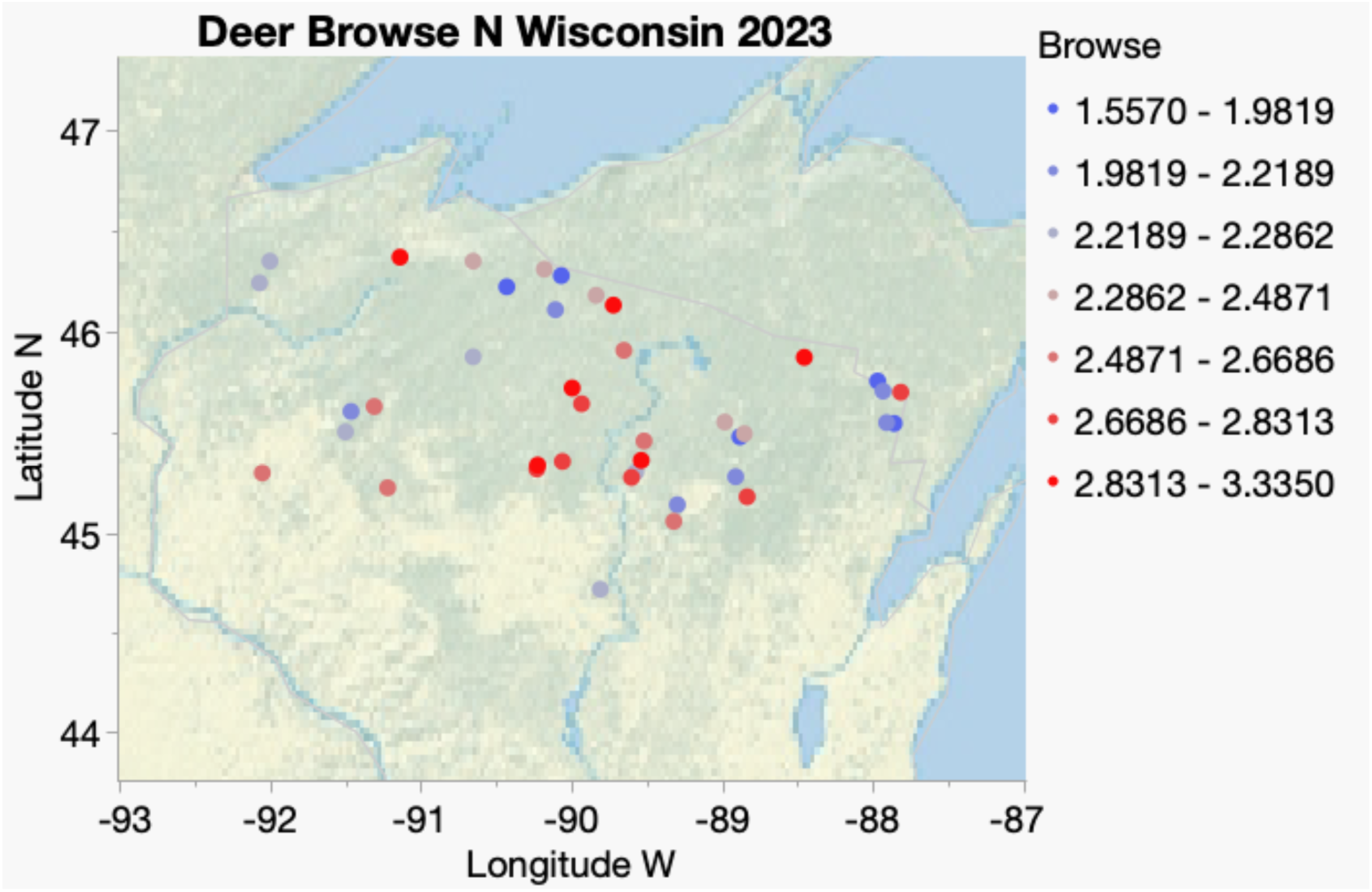
Map showing variation in the intensity of deer browse pressure in N Wiscsonsin. Browse (the converse of twig age, Eqn. 3) appears to show geographic structure with hotspots (red) and cool spots (blue).

## Discussion

Deer populations have increased across many temperate regions bringing high impacts on plant and animal diversity and tree regeneration. Developing and applying methods to monitor deer impacts across large regions and over time would greatly enhance our awareness of these trends and our ability to manage forests and wildlife more sustainably. Although many methods exist to monitor deer populations and impacts (see *Introduction*), there is no consensus regarding which method(s) are most reliable and useful. This partly reflects that few methods have been rigorously tested and compared to other methods (exception: Sample et al., 2023). This has impeded progress in determining which method(s) are the most accurate and efficient. Additional collaborative efforts are thus needed to test alternative monitoring methods so that we can standardize and assemble the tools we need to build reliable monitoring programs.

To be most useful, indicators should be valid, appropriate, reliable, accurate,, and efficient. The goals laid out in the introduction address these criteria. To test validity (goal 1), I compared twig ages on woody seedlings and saplings in- vs. outside the fenced exclosure at HMC. These results confirm that deer markedly reduce twig ages within the ‘molar zone’ (20-200cm) in all species measured (Fig. 2, Table 1, SI Fig. S4). Species, height, and other factors further affect twig ages, but deer (underestimated by the Exclosure effect) predominate with extraordinary statistical power (F=612 vs. 24 for next most important factor, Table1; F=796 vs. 52 for the next, Table 2). These results confirm that twig ages provide a valid and appropriate indicator for deer browse.

Comparing twig ages to seedling heights as indicators of browsing was goal 2. Heights are key to several monitoring schemes including forest regeneration metrics used by Wisconsin (Wisconsin DNR, 2024) and the U.S. Forest Service (McWilliams 2015); T. Rawinski’s “10-tallest” method (Rawinski, 2014); and the Assessing Vegetation Impacts of Deer (AVID) protocol (Russell & Desprez, 2022; Quirion & Blossey, 2023). Although heights respond to deer at HMC, these responses were muted and complex relative to the clear and consistent signals from twig age (e.g., a height F-value ∼1% that of twig age with variable effects across species, Table 1). Thus, it is unwise to rely on snapshot surveys of height as a browse indicator. Marking plants to monitor seedling heights over time (as in the sentinel oak method) provides a more reliable indicator but requires considerably more time and effort. Obtaining twig age snapshot data is far more efficient and generates seedling heights as a by-product.

Twig ages also provided a reliable way to track changes in browsing over time (goal 3) in the repeated surveys at HMC. Despite multiple 2-way interactions (Table 2, SI Fig. S1), year was second only to deer in predicting variation in twig age with the Year x Exclosure interaction showing increased deer impacts at HMC. Results at HMC also made clear how mean twig ages vary over species in ways that can also vary over time. Shifts in deer browse impacts over time and species are of particular interest to managers charged with sustaining both wildlife populations and successful regeneration. The fact that all three main effects and all their interactions were highly significant demonstrate how statistically well-behaved twig age data are and their power for tracking how tree seedlings respond to browsing in a nuanced way. In sum, the twig age method, first developed at HMC, provides a sensitive indicator of how deer impacts vary over space (in vs. outside the exclosure), species, and time. Most notably, it provides “real-time” indicators for how species respond to changes in deer browsing.

### Height effects

Seedling height affects the apparency of seedlings to deer and how convenient it is for deer to feed on them affecting browsing rates in ways that themselves indicate herbivory (goal 4). Recording heights should thus remain part of the twig age protocol. Twig ages responded to seedling heights at HMC in various ways depending on species and exposure to deer. At the dispersed S and N Wisconsin sites (and in *Acer pensylvanica*, *A. rubrum* and *Q. rubra* at HMC in 2022), species showed more browsing on seedlings of intermediate height. This ‘bite out of the middle’ likely reflects how deer find it more convenient to browse on twigs at head height. Given these effects, controlling for seedling height when analyzing differences in twig age among species, sites, and years tends to reduce bias and increase precision. When deer respond differently to seedling height depending on species, we should also control for height x species interactions.

### Assessing differences among species and sites

Any reliable indicator of herbivory should be sensitive to differences among species (reflecting differential palatability) and sites (reflecting spatial variation in browsing – goal 5). Sampling twig ages on multiple species in rough proportion to their natural occurrence across the nine S Wisconsin and 40 N Wisconsin sites allowed me to explore the sensitivity of twig age to both site and species effects. Mean twig ages varied widely and the customarily small standard errors yielded high power for discriminating differences among both species and sites (Fig. 3). Given the other results presented here, I interpret these differences as accurate depictions of browse impacts vary among species and sites. Nevertheless, it would be wise to confirm this using additional sites especially where deer enclosures or exclosures are available. Such tests might also confirm whether twig age metrics like ATA1 and ATA2 work most reliably for judging relative impacts on species and sites (ranks) or whether they represent interval-level data capable of providing proportional comparisons.

Interestingly, mean twig ages spanned almost similar ranges among sites in S and N Wisconsin. Data from both regions also yielded similar levels of sensitivity for assessing differences in browse pressure among sites (F=20.4 and 21.2 in S and N Wisconsin, respectively). Including seedling height effects increased this sensitivity (to F=22.2 and 23.4, respectively, SI Table S4). Differences in twig age among taxa, however, were much greater in N than in S Wisconsin (F=31.8 vs. 3.1, SI Table S3). This partly reflects the high sensitivity to browsing (very low mean twig ages) in aspen (*Populus tremuloides*) in N Wisconsin.

### Adjusting twig age data for species effects

Because species differ in growth rates and palatability, it is important to adjust for species effects when comparing sites and tracking browse impacts over time (goal 6). Fortunately, twig age differences among taxa were remarkably consistent across sites, reflecting a transitive pattern of browse susceptibility. This allows us to adjust for species effects to obtain more reliable twig age indicators (goal 7). The lack of substantial taxon x site interactions in S and N Wisconsin confirmed transitivity, supporting this approach. This led to two adjusted measures: ATA1 equivalent to adjusted least-square site means derived from 2-way ANOVAs, and ATA2 which adjusts each taxon’s mean twig age using a ratio indicating how much more palatable (browsed) it is relative to sugar maple. Both adjusted indicators account for differences in how heavily deer browse different species, mirroring the deer preferences in Fig. 3. Usefully, ATA1 and ATA2 yielded almost identical results in N Wisconsin, suggesting that either metric correctly adjusts twig age data to account for the species present at a site. Estimates based on mean twig age or just sugar maple TA at a site, however, were somewhat different (SI Fig. S6), reinforcing the value of using ATA1 or ATA2. Given the extensive sampling across 40 sites, the SM preference ratios provided here should prove useful for sites across the upper Midwest (though this assumption should be checked). Either metric also enhances sample sizes by allowing field workers to sample all suitable seedlings present at a site while adjusting for differences in species composition. Sampling seedlings in proportion to their abundance further allows one to estimate overall site palatability, a metric also worth tracking over time.

Finally, I tested whether sites experiencing more browsing now (smaller mean twig ages) show signs of having shifted in composition to have fewer palatable species. Instead, sites with more browsing showed higher fractions of palatable species (SI Fig. S7). This suggests simply that sites with many palatable twigs are browsed more, showing shorter twig ages, rather than any shift in community composition following browsing. The fact that ATA1 and ATA2 did not show any relationship to site palatability reinforces how these metrics provide more reliable ways to compare sites while avoiding biases due to differences in species composition.

### Mapping browse

Mapping adjusted twig ages for the 40 N Wisconsin sites revealed geographic variation with some structure. Because twig ages respond quickly to changing browse levels, monitoring would quickly reveal whether this structure is stable or how it might shift in response to management. Maps of browse pressure are thus useful, both for generating hypotheses about which factors drive variation in deer browse and for assessing how quickly management efforts can affect browse impacts. Knowing which factors are acting and how quickly would enhance our ability to make more informed management decisions.

### Statistical properties

Twig age estimators are statistically well behaved with compact standard errors (SE), high sensitivity to differences among species and sites, and infrequent higher-order interactions. Increased sample sizes always reduce standard error estimates, but these improvements were marginal beyond 120 twigs (Fig. 5). Such estimates from N Wisconsin had remarkably small SEs (∼0.05 years for means of ∼2 years). Adjusted mean twig age SE’s appeared to decline less at higher sample sizes than those for simple means suggesting that adjusting for species effects may also provide more honest estimates of error variation. It will be useful to compare these SE’s to those for competing indicators of local browse at matched sites in future research.

### Efficiency

The twig age method is highly efficient to implement. Once trained to recognize and count terminal bud scale scars, it is quick to age twigs. Recording spoken data in the field leaves hands free for measuring heights and examining twigs. On average, it took 45 minutes to score 106 seedlings at the N Wisconsin sites using vocal data entry. Similar times were then needed to enter and check the data (as with paper data), but this could be accelerated using voice-to-text translation software to automate data entry into a spreadsheet (Toczydlowski, 2017). The twig age method appears to be more efficient than methods that require marking and remeasuring plots or plants, allowing many more sites to be assayed. Within sites, sampling seedlings of all taxa in proportion to their abundance further speeds data acquisition while providing information on overall seedling composition and palatability.

### Conclusions

The twig ages provide a consistent and sensitive way to measure and track deer impacts on regenerating seedlings and saplings over time and among sites. Being based on discrete data with five categories improves consistency within and among observers. The additional tests and results presented confirm how twig age data genuinely reflect deer impacts. These data also allow us to rank the relative palatability among species on a transitive scale. This scale, in turn, allows us to adjust twig age data for different palatabilities among species using either 2-way ANOVA to adjust site means (ATA1) or browsing preference ratios to adjust twig ages to the sugar maple standard (ATA2). These extensions enhance the versatility and generality of the twig age method providing a sensitive and consistent way to compare deer impacts among sites and over time. Twig age data are also nicely cumulative, allowing us to track changes in browse pressure in real time among sites and map those differences across landscapes. These tools can therefore be used to decode the factors affecting local deer impacts and how impacts respond to changes in forest and wildlife management.

In comparing several alternative browse indicators in Indiana, Sample et al. (2023) found browsing intensity and deer density in Indiana to track twig age data most closely. They concluded it was “the most efficient and effective index to monitor browse intensity in agriculturally dominated landscapes, such as those of the Central Hardwood Forest Region.” The results presented here suggest this conclusion extends to upper Great Lakes landscapes. The twig age method thus merits further use and testing to determine whether it can serve as the basis for the efficient and effective network we need to monitor deer impacts and how these respond to shifts in forest and wildlife management.

## Acknowledgements

I thank J. Witt and S. Johnson for helping develop the initial twig age method. The Huron Mountain Wildlife Foundation supported this work. W.S. Alverson, and J. Curteau.provived useful comments on the manuscript. C. Allen-Savietta suggested particular analyses.

## Data Availability

All data are available at Dryad for use by the journal and reviewers at: http://datadryad.org/stash/share/KaySv4IeQJTmgAqacnc-czrgMLGAgEQAyhuo04ew8W4. These will be shared on Dryad upon publication at DOI: 10.5061/dryad.8gtht76z5.

## Supplementary Information

**Supplementary Table S1.** List of study sites, year(s) surveyed, and sample sizes showing the number of seedlings/saplings measured for twig age at each site, by genus.

| Location | Survey<br>Year | Acer | Carya | Fraxinus | Ostrya | Quercus | Other | All<br>Species |
| --- | --- | --- | --- | --- | --- | --- | --- | --- |
| **Huron Mountain Club** | 2015 | 202 |  |  |  |  |  | 202 |
|  | 2016 | 87 |  |  |  |  |  | 87 |
|  | 2017 | 303 |  |  |  |  |  | 303 |
|  | 2019 | 288 |  |  |  |  |  | 288 |
|  | 2022 | 181 | 34 |  |  | 10 | 22 | 247 |
| **S Wisconsin Sites** |  |  |  |  |  |  |  |  |
| Abraham's Woods | 2019 | 32 |  |  |  |  |  | 32 |
| Baxter's Hollow | 2019 | 53 |  | 51 |  | 5 |  | 109 |
| Botany Farm | 2019 | 54 | 27 | 54 |  |  |  | 135 |
| Devil's Lake, Burma | 2019 | 60 | 9 | 51 |  | 7 |  | 127 |
| Devil's L. Oak Forest | 2019 | 56 | 9 | 65 |  |  |  | 130 |
| Happy Hill Glades | 2019 | 53 | 52 | 25 |  | 44 |  | 174 |
| Leopold Reserve | 2019 | 82 |  |  |  | 140 |  | 222 |
| McGilvra | 2019 | 50 |  | 51 |  |  |  | 101 |
| UW Arboretum | 2019 | 79 |  |  |  |  |  | 79 |
| **N Wisconsin Sites** |  |  |  |  |  |  |  |  |
| Arbor Vitae | 2023 | 6 |  |  | 16 | 97 | 12 | 131 |
| Bergen | 2023 | 2 |  | 39 |  | 11 | 62 | 114 |
| Berry La. | 2023 | 77 |  | 41 | 1 | 1 |  | 120 |
| Bibon | 2023 | 47 |  | 93 | 6 | 32 | 16 | 194 |
| Blossey 1 | 2023 | 20 |  | 11 |  |  | 12 | 43 |
| Blossey 2 | 2023 | 0 |  | 27 |  | 5 | 28 | 60 |
| Blossey 3 | 2023 | 37 |  |  | 18 |  |  | 55 |
| Blossey 4 | 2023 | 50 |  | 4 | 1 | 3 | 1 | 59 |
| Blossey 5 | 2023 | 13 |  | 16 |  |  | 21 | 50 |
| Blossey 6 | 2023 | 11 |  | 9 |  | 15 | 14 | 49 |
| Blossey 7 | 2023 | 2 |  | 20 |  | 29 | 5 | 56 |
| Blossey 8 | 2023 | 5 |  |  | 20 |  | 7 | 32 |
| Blossey 9 | 2023 | 5 |  | 31 | 13 | 5 | 2 | 56 |
| Blossey 10 | 2023 | 44 |  | 11 |  |  | 1 | 56 |
| Blossey 11 | 2023 | 44 |  | 12 |  |  | 6 | 62 |
| Boulder Jct. | 2023 | 2 |  |  | 57 | 47 | 12 | 118 |
| Carey | 2023 | 40 |  | 1 | 39 | 5 | 15 | 100 |
| Copper Falls | 2023 | 78 |  | 42 |  |  | 2 | 122 |
| Dairyland | 2023 | 47 |  | 26 | 4 | 32 | 38 | 147 |
| Gordon | 2023 | 70 |  | 12 | 10 | 15 | 22 | 129 |
| Harrison | 2023 | 5 |  |  | 2 | 37 | 75 | 119 |
| Holcomb-Birch Ck. | 2023 | 13 |  | 50 | 6 | 61 |  | 130 |
| Ice Age Trail | 2023 | 76 | 5 | 52 | 7 | 2 | 6 | 148 |
| Jacobs | 2023 | 100 |  |  | 6 |  | 10 | 116 |
| Lake | 2023 | 18 |  | 77 | 5 | 11 | 18 | 129 |
| Lynne N | 2023 | 103 |  | 72 | 6 |  | 10 | 191 |
| Lynne S | 2023 | 91 |  |  | 23 | 1 | 5 | 120 |
| Mercer | 2023 | 96 |  |  | 2 |  |  | 98 |
| Merrill | 2023 | 43 |  | 25 | 30 | 12 | 25 | 135 |
| Merrill N Loop | 2023 | 45 |  | 2 | 28 | 17 | 19 | 111 |
| Murry | 2023 | 6 | 7 | 55 | 20 |  | 11 | 99 |
| Oma | 2023 | 102 |  | 2 | 1 |  | 4 | 109 |
| Rib Lake 1 | 2023 | 48 |  | 81 | 12 | 32 | 24 | 197 |
| Rib Lake 2 | 2023 | 47 |  | 28 | 10 | 19 | 32 | 136 |
| Russell | 2023 | 21 |  | 37 | 2 | 72 | 5 | 137 |
| Sherman | 2023 | 68 |  | 4 | 13 | 2 | 25 | 112 |
| Turtle Lake | 2023 | 9 | 1 | 42 | 15 |  | 39 | 106 |
| Wilkinson | 2023 | 25 | 4 | 38 | 14 | 8 | 21 | 110 |
| Wilson | 2023 | 9 |  | 43 | 4 | 47 | 18 | 121 |
| Winchester | 2023 | 72 |  | 4 | 26 |  | 9 | 111 |
| **Totals** |  | **3177** | **148** | **1304** | **417** | **824** | **654** | **6524** |

**Supplementary Table S2.** Results of the GLM analysis at HMC which assessed variation in 2022 twig ages among six woody taxa and the fenced exclosure (deer effects). The one significant 2-way interactive effect is shown but species x exclosure was NS, indicating that species twig ages responded consistently to deer browsing. R^2^=0.52, N=247 (cf. Fig. 3). The equivalent analysis for seedling height had R^2^=0.19 and an Exclosure F=6.38.

| **Source** | **DF** | **S.S.** | **F Ratio** | **Prob > F** |
| --- | --- | --- | --- | --- |
| **Exclosure** | 1 | 126.54 | 183.81 | <0.0001 |
| **Species** | 5 | 41.06 | 11.93 | <0.0001 |
| **Height** | 1 | 12.19 | 17.70 | <0.0001 |
| **Species\*Height** | 5 | 18.42 | 5.35 | 0.0001 |

**SI Table S3.** Results of GLM analyses (2-way ANOVA’s) of variation in twig age across the 8 taxa and 9 sites of S Wisconsin (a, N=1109, r2=0.22) and the 10 taxa and 40 sites of N Wisconsin (b, N=4110, r2=0.26).

| Source | DF | Sum of Squares | F Ratio | Prob > F |
| --- | --- | --- | --- | --- |
| Taxon | 7 | 14.48389 | 3.0865 | 0.0032* |
| Site | 8 | 109.59443 | 20.4349 | <.0001* |

b)
| Source | DF | Sum of Squares | F Ratio | Prob > F |
| --- | --- | --- | --- | --- |
| Taxon | 9 | 188.03647 | 31.7869 | <.0001* |
| Locale | 39 | 544.44851 | 21.2393 | <.0001* |

**SI Table S4.** Effects of seedling heights on twig age among the 9 taxa and 10 sites of S Wisconsin (a, r2=0.293, N=1109) and 10 taxa and 40 sites of N Wisconsin (b, r2=0.297, N=4108). Note the strong quadratic effects of height, reflecting preferential deer browsing at intermediate seedling heights (Fig. 5). The Taxon x Height interaction was only significant in N Wisconsin.

| Source | DF | Sum of Squares | F Ratio | Prob > F |
| --- | --- | --- | --- | --- |
| Site | 8 | 110.42025 | 22.2092 | <.0001* |
| Taxon | 7 | 16.34531 | 3.7572 | 0.0005* |
| Height * Height | 1 | 4.19354 | 6.7477 | 0.0095* |
| Height * Taxon | 7 | 17.62845 | 4.0522 | 0.0002* |
| Height | 1 | 1.26660 | 2.0381 | 0.1537 |

b) N Wisconsin
| Source | DF | Sum of Squares | F Ratio | Prob > F |
| --- | --- | --- | --- | --- |
| Locale | 39 | 568.46161 | 23.4402 | <.0001* |
| Taxon | 9 | 166.41915 | 29.7361 | <.0001* |
| Height | 1 | 24.78159 | 39.8523 | <.0001* |
| Height * Height | 1 | 94.70330 | 152.2962 | <.0001* |
| Height * Taxon | 9 | 37.12903 | 6.6343 | <.0001* |

**SI Table S5.** Results of a general linear model relating mean twig age in ten deciduous taxa to the mean Sugar Maple twig age at the same site across all 40 sites in N Wisconsin. The model accounts for 35.1% of the total variance with F=4.29 (p<0.0001) and N=171.

| Source | DF | Sum of Squares | F Ratio | Prob > F |
| --- | --- | --- | --- | --- |
| Taxon | 9 | 14.807394 | 5.8582 | <.0001* |
| SM Twig Age | 1 | 3.001086 | 10.6858 | 0.0013* |
| SM Twig Age*Taxon | 9 | 2.121425 | 0.8393 | 0.5811 |

**SI Figure S1.**
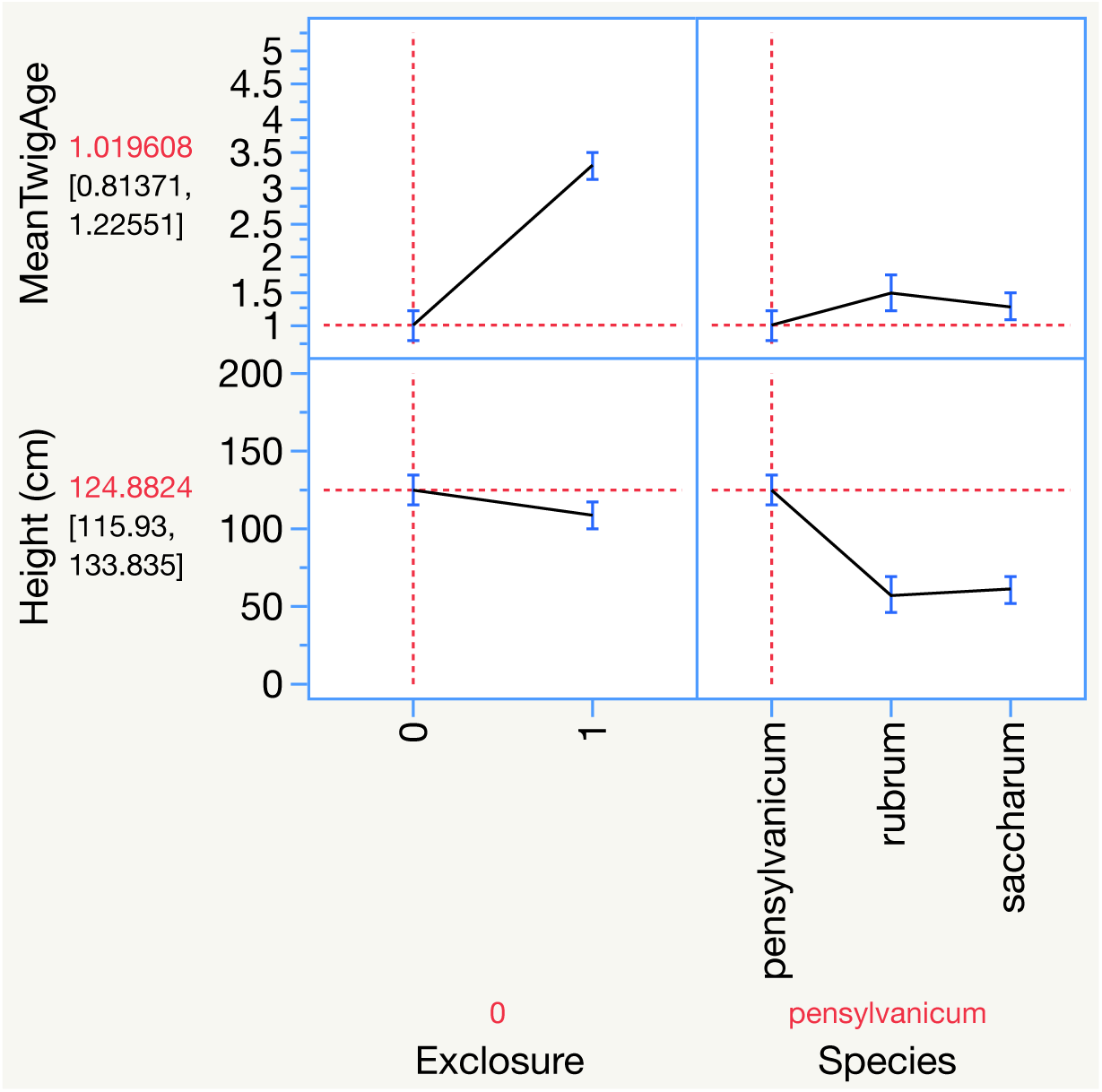
Profile plots showing the main effects of exclosure and species on mean twig age (above) and seedling height (below) on three species of maple (*Acer*) growing at Huron Mountain Club. Least square means and standard errors are plotted, adjuisted for other effcts. Note twig age is far more responsive to deer than seedling height (exclosure effect - left) while being far *less* responsive to species (right).

**SI Figure S2.**
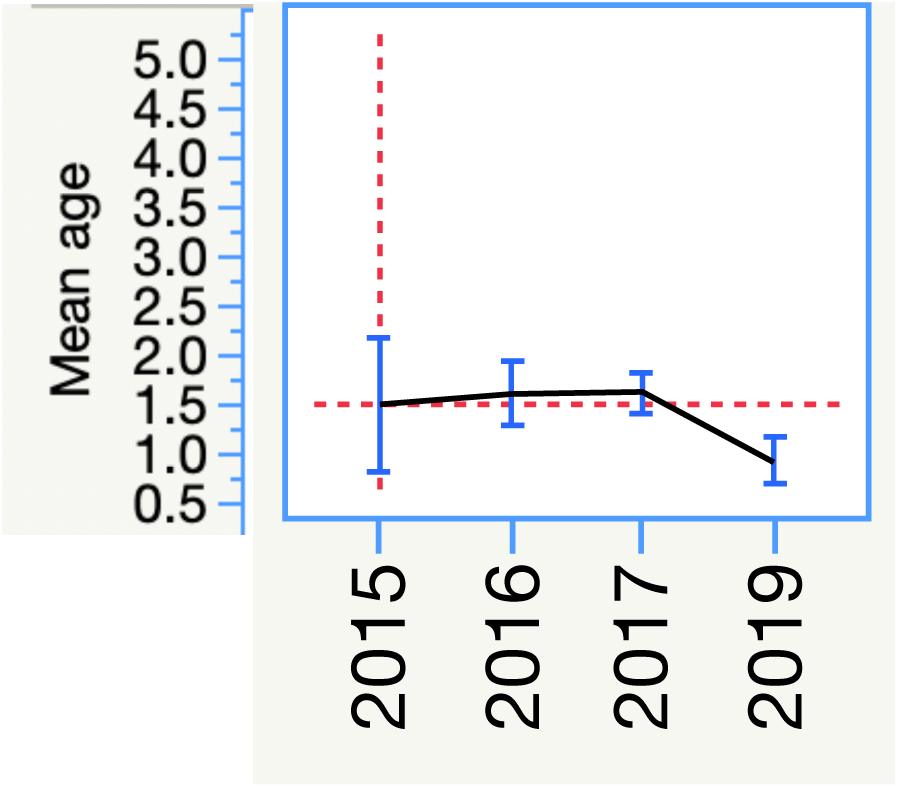
Profile plot showing the main effect of year in analyses of mean twig age across three species of maple and four years at Huron Mountain Club. Corresponds to results in Table 2.

**SI Figure S3.**
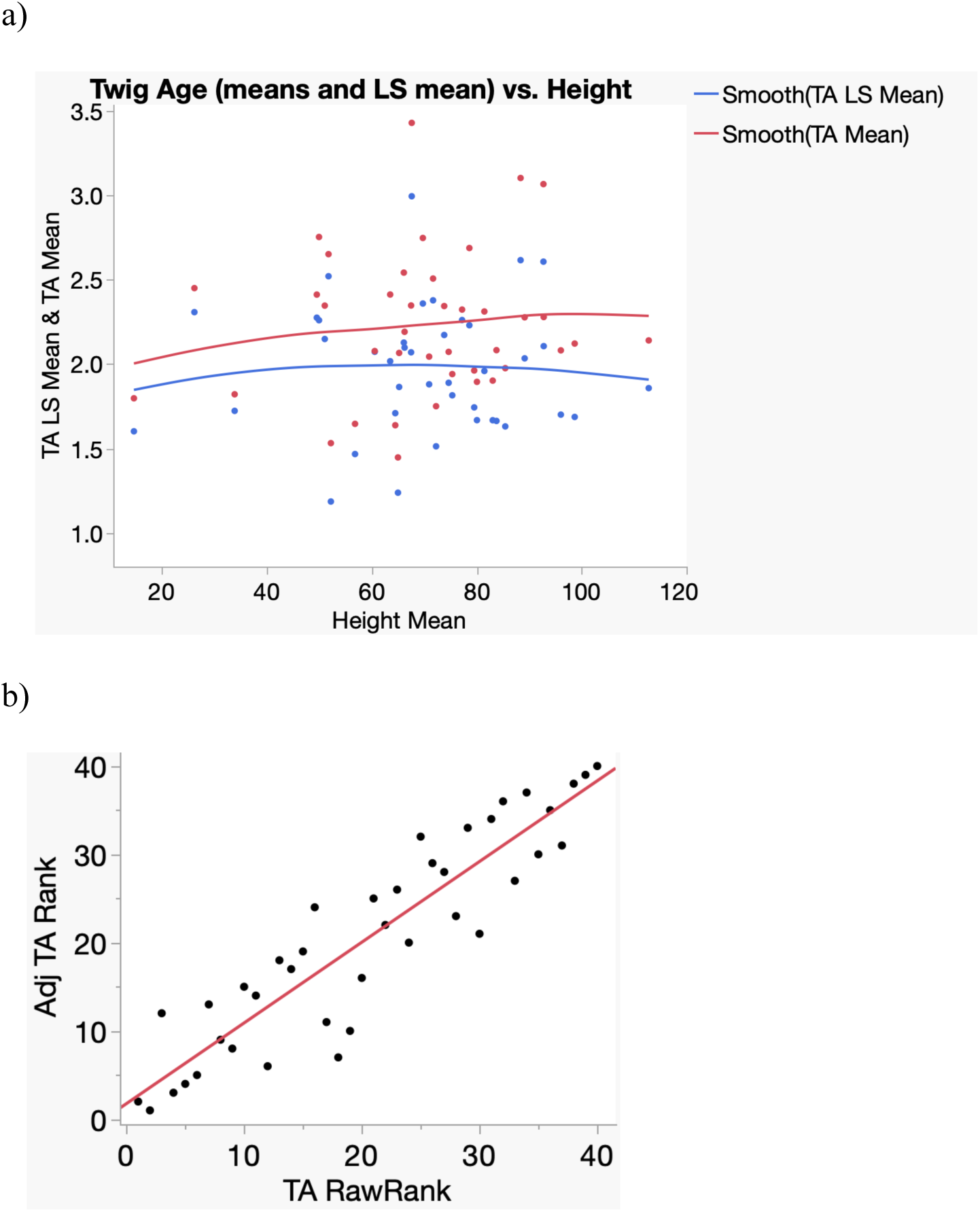
Effects of adjusting mean site twig ages for species effects among the 40 N Wisconsin sites. Adjusting for species effects slightly reduced estimated mean twig ages, as expected if fewer palatable seedlings persist at more heavily browsed sites (a). Adjusting for species effects slightly affected how the 40 sites were ranked for deer impacts (b). Spearman rank correlation: ρ=0.91.

**SI Fig. S4.**
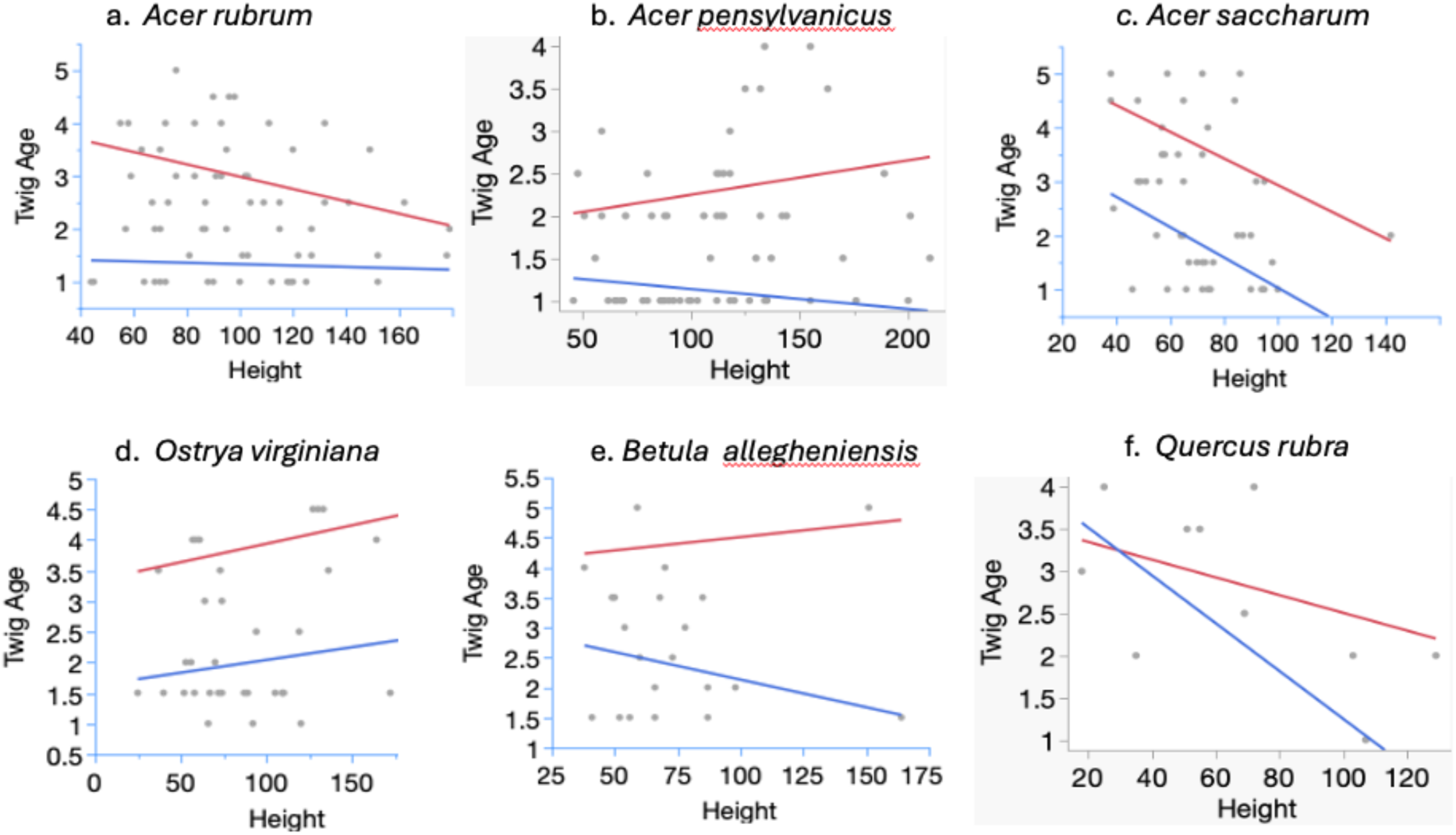
Twig age responses to seedling height and the deer exclosure varied among six species sampled at HMC in 2022. Red lines fit seedlings inside the exclosure while blue lines fit seedlings outside the fence, reflecting strong effects of deer on both mean twig age and how twig ages respond to height.

**SI Figure S5.**
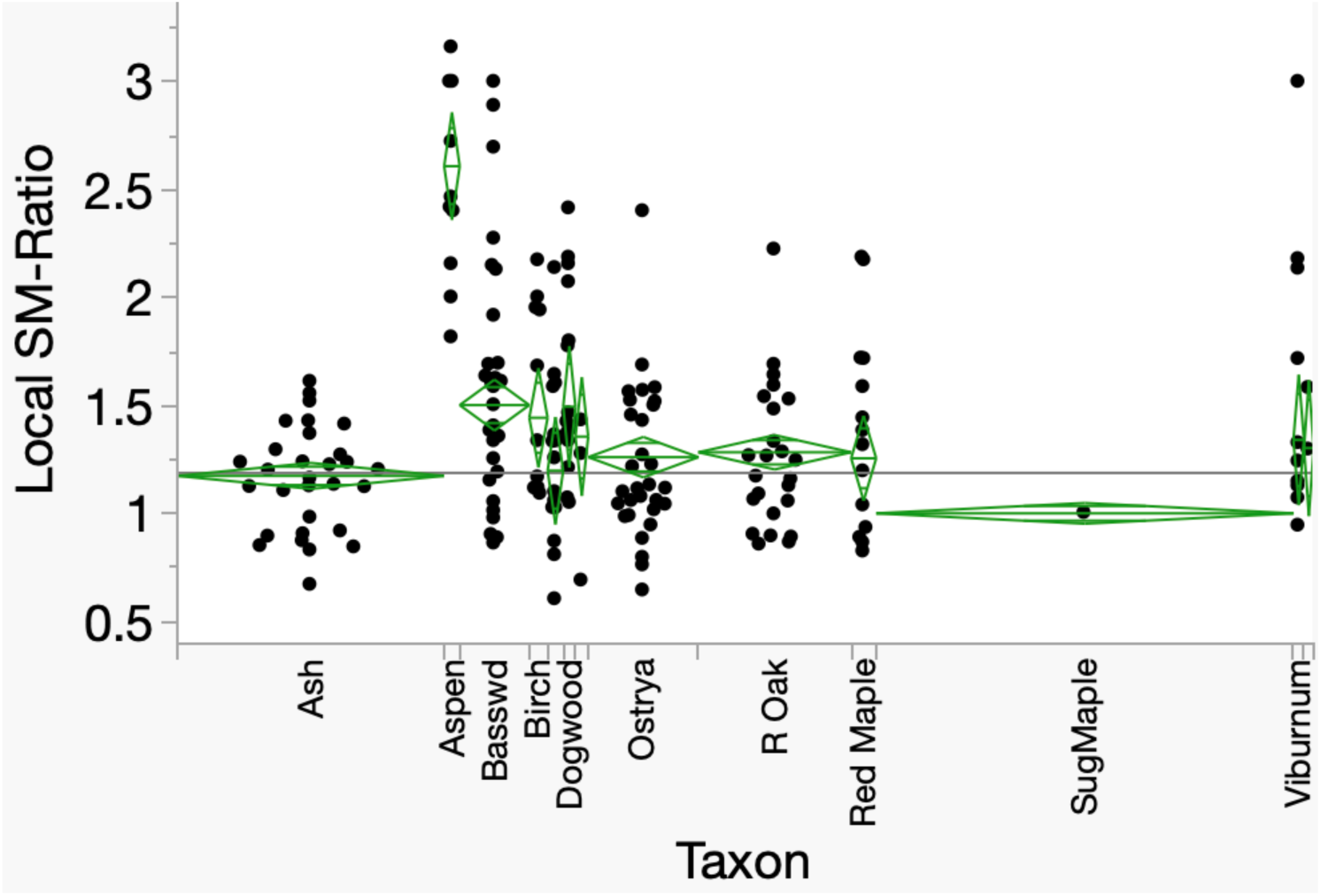
One-way ANOVA of variation in local twig ratios (sugar maple to other taxa). Taxon accounts for 52% of this variance (F=19.5, p<0.0001). Width of columns reflects sample sizes for each taxon.

**SI Figure S6.**
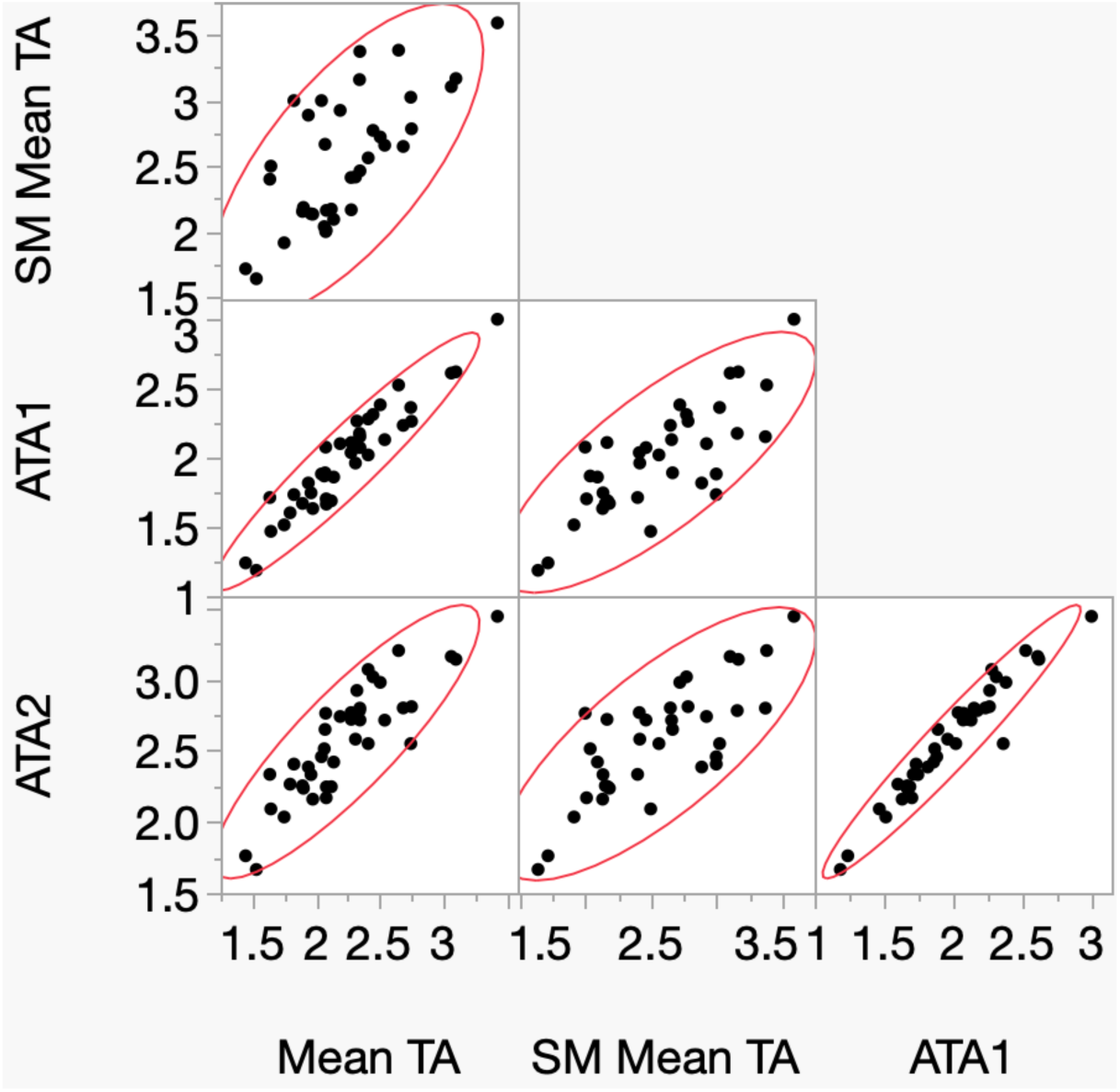
Relationships among four estimators of deer browse based on twig age: mean twig age at a site (Mean TA); the least square mean twig age, corrected for species effects (ATA1); mean twig age only for sugar maple (SM Mean TA); and all species adjusted to the sugar maple standard twig age (ATA2). ATA1 and ATA2 correct for species differences in palatability in different ways but are essentially identical (r=0.97).

**SI Figure S7.**
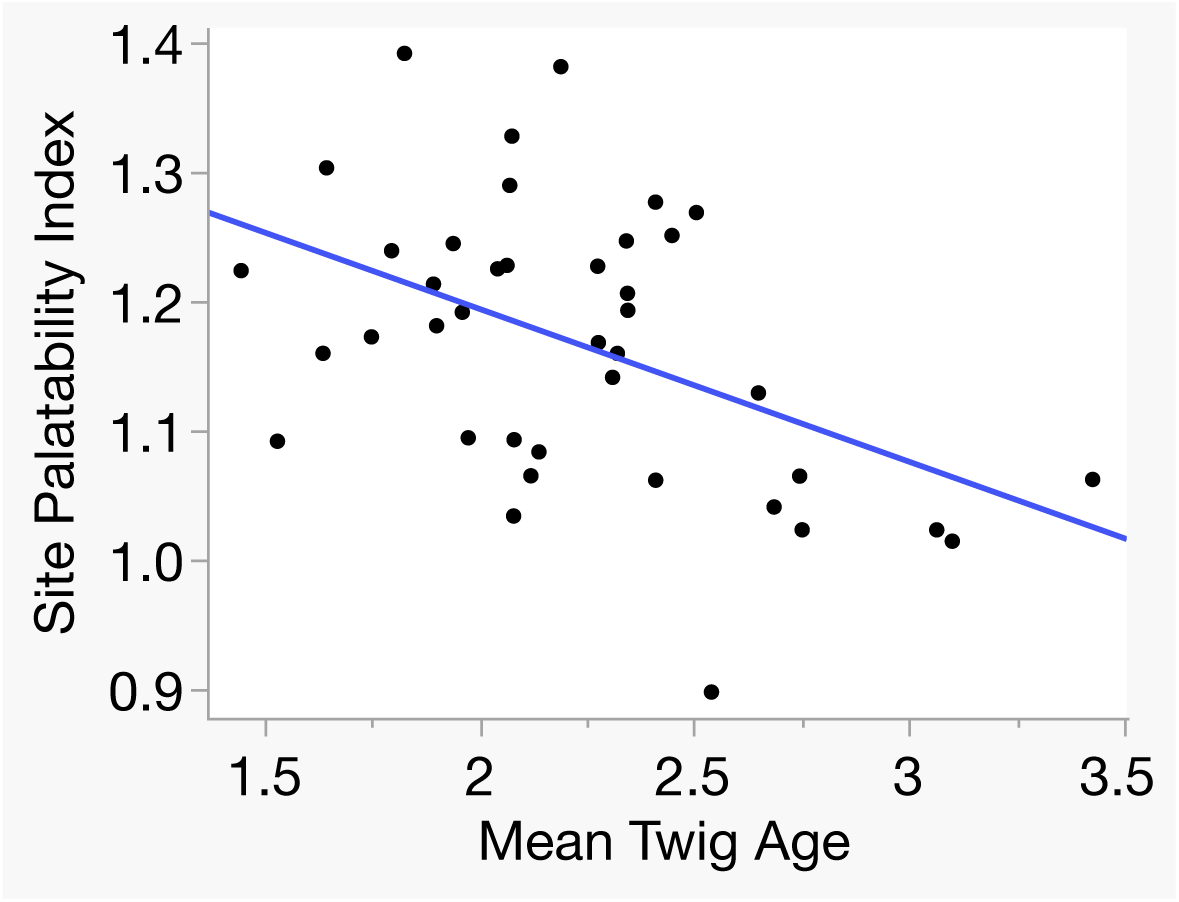
The mean palatability of seedlings present declines as mean twig age increases across sites in N Wisconsin. R=-0.47, p = 0.0025. This likely reflects the fact that twig ages are higher in less palatable species which, when common, cause the Site Palatability Index to fall. Adjusted Twig Age and the Browse index showed no relation to site palatability. Sherman Corners had the lowest site palatability index (0.9) but intermediate twig age.

